# Non-autonomous PMK-1/p38 MAPK signaling ensures homeostatic downregulation of germline stem cell proliferation in *C. elegans*

**DOI:** 10.1101/2025.09.24.678176

**Authors:** Alexandre Clouet, Matthieu Valet, Benjamin Dufour, Janina Rieger, Xavier Lechasseur, Aurelia Thérèse Condomat Zoumabo, Lloyd Venceslas Fotso Dzuna, Alexane Murray, Jichao Deng, Pier-Olivier Martel, Patrick Narbonne

**Author notes:** **Corresponding author’s email address**.

## Abstract

Stem cell underproliferation leads to tissue loss while over proliferation favors hyperplasia and tumorigenesis. Stem and progenitor proliferation rates are therefore usually coupled to the needs for their differentiated cellular progenies in the tissue(s) they serve. We used the *C. elegans* hermaphrodite reproductive system to understand how germline stem cell (GSC) proliferation is linked to sperm availability and oocyte usage. We developed a forward genetic screening method to identify genes required for the homeostatic downregulation of GSC proliferation when oocytes are not used, where germline homeostasis defective (*ghd*) mutants grow benign differentiated germline tumors. We provide evidence that the TIR-1/SARM1 metabolic sensor works together with the NIPI-3/TRIB1 pseudokinase to activate a canonical PMK-1/p38 MAPK signaling cascade in the somatic gonad, shut down the extracellular regulated kinase MPK-1/ERK, and couple GSC proliferation to oocyte needs. Similar multi-tissue signaling modules could underlie homeostatic stem cell regulation and prevent benign tumorigenesis in humans.

## Introduction

Stem cells (SCs) are essential throughout the life of organisms. They are required to generate functional tissues and organs during development and to ensure a healthy cellular turnover during adulthood. However, as each of their division comes with the risk of introducing replication and segregation errors, such that an increased number of SC divisions predisposes to cancer^1,2^, SC proliferation rates must be tightly regulated to prevent any unnecessary division.

SC proliferation may be influenced by systemic elements, such as hormones and growth factors. Notably, nutrient uptake sustains SC proliferation through activating the conserved insulin/insulin-like growth factor 1 signaling (IIS) pathway^3^. Tissue-specific homeostatic regulatory mechanisms can further link stem/progenitor cell proliferation to various cues, including a tissue’s need for their differentiated cellular progeny. For instance, abundant differentiated progeny sends a paracrine signal through FGF18 and BMP6 to promote bulge SC quiescence in murine hair follicles^4^. Negative feedback loops required for downregulation of immature cell proliferation were also found in invertebrates. In *Drosophila* larvae, a multi-tissue signal is initiated by the hematopoietic progenitor niche and integrated by differentiated hemocytes, resulting in adenosine deaminase Adgf-A secretion to inhibit the extracellular mitogenic function of adenosine by converting it into inosine^5^. Furthermore, in the adult midgut, damaged, infected or stressed enterocytes secrete cytokines that promote intestinal SC proliferation through the Jak/Stat pathway^6^.

Another example has been observed in *C. elegans* hermaphrodites. During the last larval stage, a limited number of spermatozoa are generated and stored in spermathecae^7^. In adults, the germline stem cell (GSC) differentiation program irreversibly switches to oocyte production, which are continually fertilized until spermathecae are emptied. When this happens, oocytes accumulate in proximal gonad arms, and homeostatic feedback represses GSC proliferation to prevent oocyte overaccumulation^8–10^. Germline feminizing mutations (*i.e. fog* and *fem* mutants)^11,12^ that prevent sperm formation trigger homeostatic downregulation of GSC proliferation in young adults, while this process can be reversed upon mating with a male^8^. Downregulation of GSC proliferation is also irreversibly triggered in oocyte maturation defective *oma-1/2* mutants in which sperm is present, but oocytes are activation incompetent^10,13^. Since arrested oocytes are not laid in this background, it creates a valuable paradigm to study the formation of benign differentiated tumors as oocytes keep accumulating in *oma-1/2* mutants that are further defective in GSC homeostasis regulation^8,10,14^.

A gain-of-function (*gf*) allele in the *C. elegans* Rat sarcoma (Ras) ortholog *let-60* prevents the homeostatic downregulation of GSC proliferation^10^, while LET-60 is a key activator of the extracellular regulated kinase (ERK) ortholog MPK-1^15,16^, an important positive regulator of GSC proliferation^10,17^. Interestingly, MPK-1 promotes GSC proliferation cell non-autonomously as transgenic rescue of the soma-specific MPK-1A isoform restores GSC proliferation in *mpk-1(Ø)* mutants^18^. Therefore, somatic MPK-1 may be inhibited by homeostatic signaling to induce GSC quiescence.

Several factors are required for homeostatic downregulation of GSC proliferation. These include the *C. elegans* orthologs of the liver kinase B1 (LKB1), the AMP-activated protein kinase α2 catalytic subunit (PRKAA2), the phosphatase and tensin homolog (PTEN), and 14-3-3: PAR-4, AAK-1, DAF-18 and PAR-5, respectively. Depletion of any of these gene products disrupt germline homeostasis and prevents the downregulation of GSC proliferation in the absence of sperm^10^. In these spermless homeostasis-defective double mutants, GSC proliferation and oocyte production proceed at a fast pace, as in the wild-type, while oocytes spontaneously mature, are ovulated and laid as endomitotic oocytes. Apart from an epistasis analysis that placed *daf-18* upstream of *mpk-1* inhibition^10^ and the direct phosphorylation link between PAR-4/LKB1 and AAK-1/AMPK^19,20^, how the components of this cryptic cascade assemble to link oocyte needs to MPK-1 activity in somatic tissues remains unclear. The identification of new germline homeostasis effectors along with their spatial requirements and targets would represent a significant step towards decrypting this mechanism.

Here, we present a forward genetic screen designed to identify new germline homeostasis defective (*ghd*) mutants. This screen highlighted a new tumor suppressor role for the *C. elegans* ortholog of Sterile Alpha and TIR motif containing 1 (SARM1) TIR-1. This led us to implicate a conserved p38 MAPK signaling module in the regulation of GSC homeostasis. We propose a model in which accumulating oocytes in the proximal somatic gonad initiate a PMK-1/p38-dependent stress-like response leading to the downregulation of somatic MPK-1/ERK activity and hence, of GSC proliferation.

## Results

### A forward genetic screen to identify germline homeostasis-defective mutants

To identify new genes required for homeostatic repression of GSC proliferation, we designed a new forward genetic screen. We used ethyl methanesulfonate (EMS)^21^ to mutagenize *oma-1/+; oma-2/+* L4 larvae and screened their *oma-1/2* F2 progeny for mutations that would cause the formation of benign differentiated germline tumors, likely owing to a disruption in GSC homeostasis regulation (Figure 1A)^10^. Mutagenized animals carried a transgene that labelled their germ membranes with green fluorescence^22^ to facilitate tumor detection under a low-magnification fluorescence microscope (Figure 1A). Over two small-scale screens covering approximatively 8000 haploid genomes, we isolated 9 independent *ghd* candidates. Whole genome sequencing of these candidates revealed that 26 genes were affected by at least two independent mutations.

**Figure 1.**
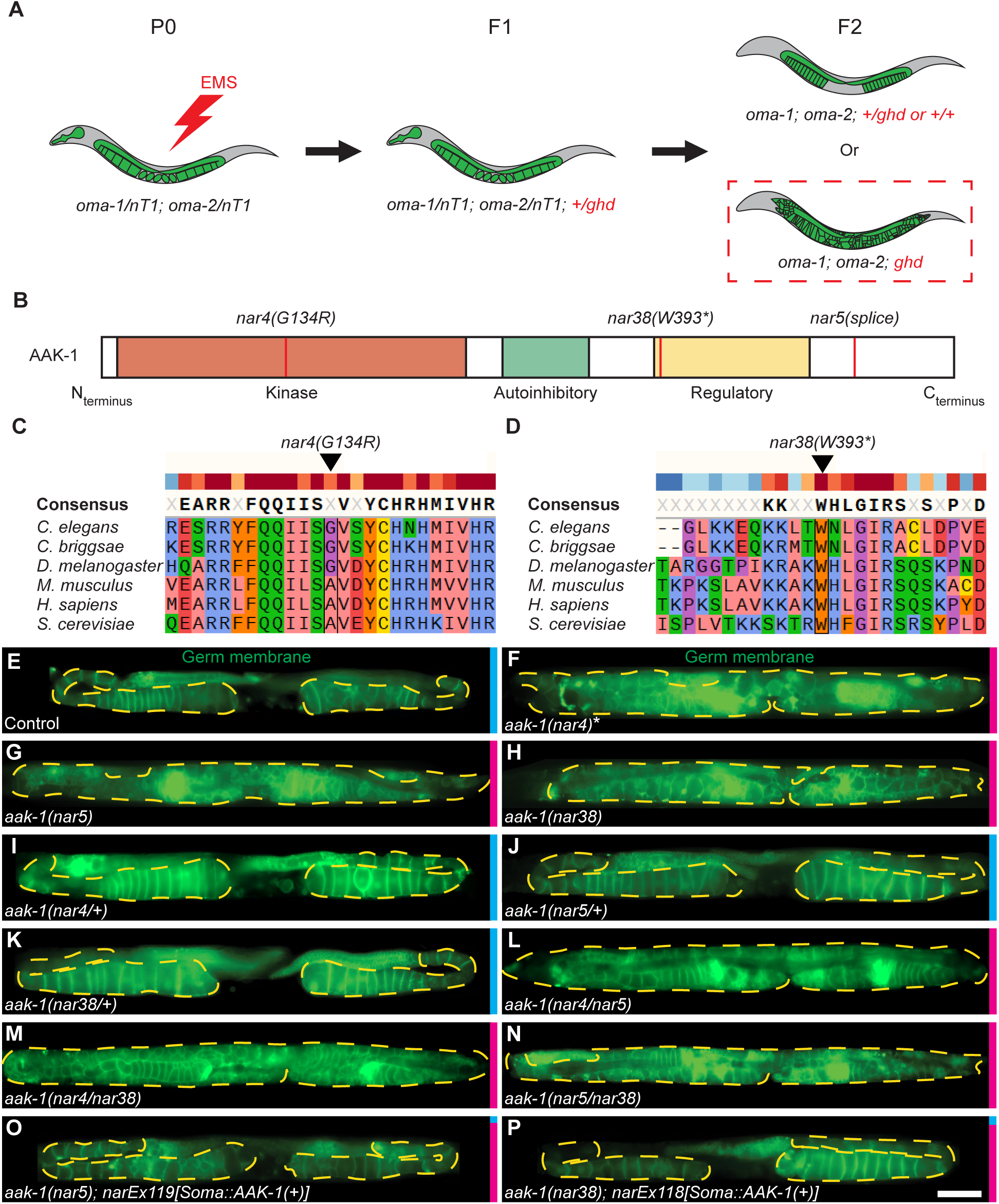
A forward genetic screen to isolate germline homeostasis-defective (*ghd*) mutants. **(A)** Schematic representation of the forward genetic screening strategy. Complete genotype of the P0 is *cpSi42[Pmex-5::mNG::PLCδ-PH::tbb-2 3’UTR + unc-119(+)]II; oma-1(zu405te33)/nT1[qIs51] IV; oma-2(te51)/nT1 V*. The *cpSi42* allele marks germ membranes with green fluorescence. The *qIs51* allele contains *Pmyo-2::GFP*, which is specifically expressed in the pharynx and marks *oma-1/+; oma-2/+* heterozygotes, but not *oma-1; oma-2* homozygotes. EMS, Ethyl Methane Sulfonate. **(B)** Schematic representation of AAK-1’s conserved functional domains. Red vertical bars, EMS-generated alleles. Asterisk, STOP codon. **(C-D)** Clustal Omega alignments of AAK-1 orthologs showing a high evolutionary conservation around the *nar4* missense (left) and *nar38* nonsense (right) alleles. **(E-P)** Representative epifluorescence micrographs of A3 hermaphrodites (20°C) of the indicated genotypes, all in the *oma-1; oma-2* background (sample sizes: 14-16-14-24-16-19-17-8-5-9-12-10). Germ membranes are marked in green by *cpSi42*. Yellow dashed lines delineate area occupied by oocytes. Anterior, left; dorsal, up. Scale bar, 100µm. A quantification of the phenotype’s penetrance is shown as a colored vertical bar on the right (Blue, Oma; Red, Ghd). All strains contain *cpSi42[Pmex-5::mNG::PLCδ-PH::tbb-2 3’UTR + unc-119(+)]*, *oma-1(zu405te33), and oma-2(te51).* *, Contains a second-site *tir-1(nar65)* mutation (see Fig. 2). (I-K) F1 cross-progeny of *oma-1/+; oma-2/+; ghd* hermaphrodites mated with control *oma-1/+; oma-2/+* males. (L-N) F1 cross progeny from reciprocal *oma-1/+; oma-2/+; ghd* crosses. (O-P) Animals were injected with a *Psur-5::GFP::AAK-1A(+)* rescue plasmid.

One of the genes mutated in three of the *ghd* candidates was *aak-1*, essential for homeostatic downregulation of GSC proliferation (Figure 1B-H)^10^. Consistent with *aak-1* mutations being responsible for the Ghd phenotype of these three candidates, they failed to complement each other (Figures 1I-N). Furthermore, transgenic somatic expression of *aak-1(+)* suppressed tumors in both *ghd-1(nar5)* and *ghd-1(nar38)* (Figures 1O-P; rescue of *ghd-1(nar4)* was not attempted due to a second-site mutation in *tir-1* as detailed below), confirming that *ghd-1* corresponded to *aak-1*. Importantly, identification of three independent *aak-1* alleles established the screen’s efficiency at isolating *ghd* mutants.

### TIR-1/SARM1 prevents the formation of differentiated germline tumors

Since genes with multiple hits in our screen were more likely to represent genuine *ghd* candidates, we fed *oma-1/2* double homozygotes with RNAi clones targeting each of them, looking for one that caused a Ghd phenotype. We found that RNAi against *tir-1*/SARM1, led to differentiated germline tumor formation in the *oma-1/2* background, and thus phenocopied the two *ghd* candidates that carried a *tir-1* mutation (Figure 2A-E, Table S1). To confirm that *tir-1* was required to prevent tumorigenesis, we examined two available predicted null *tir-1* deletion alleles: *tir-1(ok2859)* and *tir-1(qd4)*^23^ in the *oma-1/2* background. As expected, both deletions induced differentiated tumorigenesis in this background (Figure 2F-G). Moreover, neither *ghd-2(nar42)* nor *ghd-2(nar65)* complemented *tir-1(ok2859)*, suggesting that *ghd-2* corresponds to *tir-1* (Figure 2H-L).

**Figure 2.**
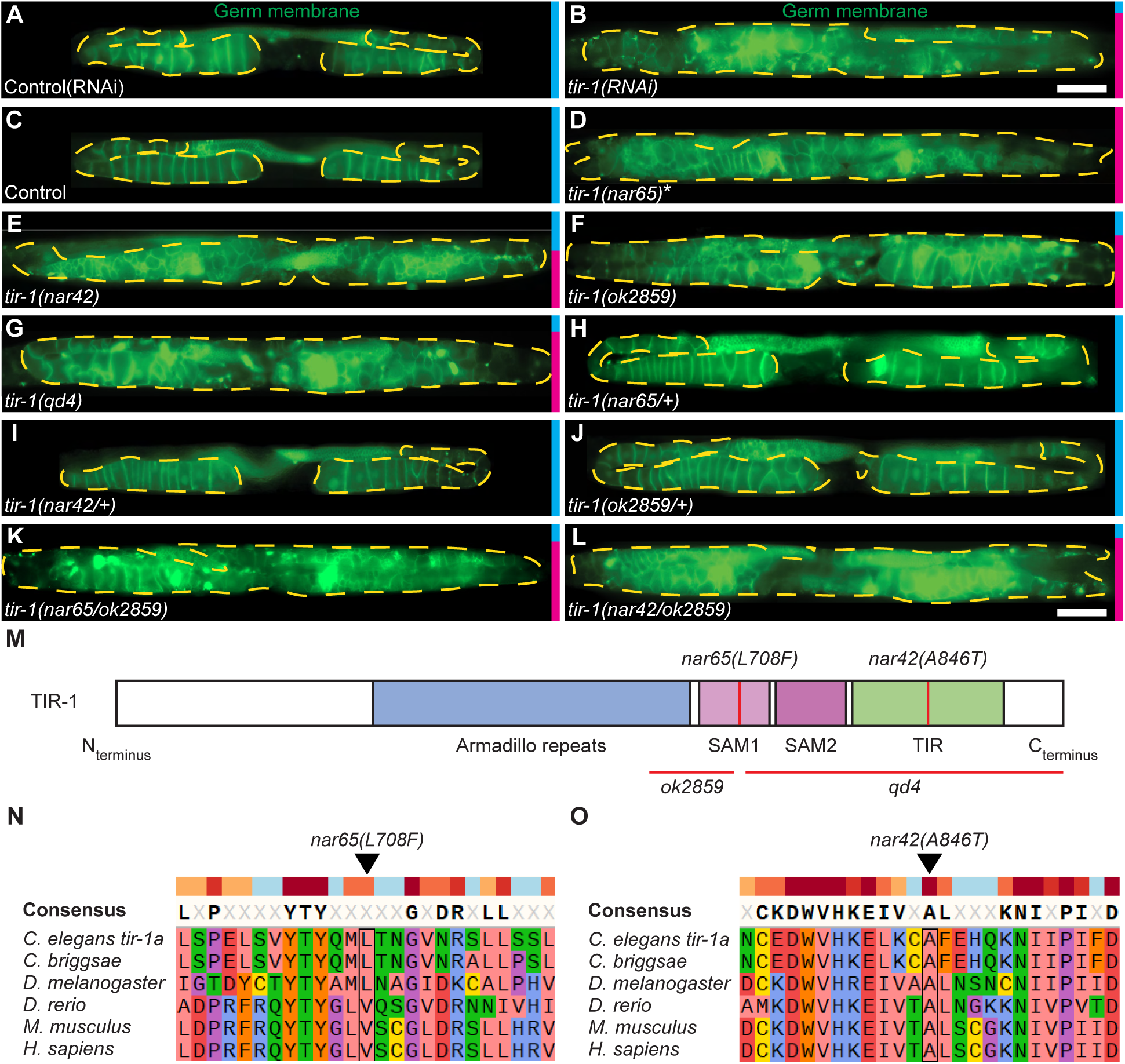
TIR-1/SARM1 prevents the formation of differentiated germline tumors. **(A-L)** Representative epifluorescence micrographs of A3 hermaphrodites (A-B: 25°C; C-L: 20°C) of the indicated genotypes, all in the *oma-1; oma-2* background (sample sizes: 12-32-14-16-22-49-12-16-21-20-11-10). Germ membranes are marked in green by *cpSi42*. Yellow dashed lines delineate area occupied by oocytes. Anterior, left; dorsal, up. Scale bar, 100µm. A quantification of the phenotype’s penetrance is shown as a colored vertical bar on the right (Blue, Oma; Red, Ghd). All strains contain *cpSi42[Pmex-5::mNG::PLCδ-PH::tbb-2 3’UTR + unc-119(+)]*, *oma-1(zu405te33), and oma-2(te51).* *, Contains a second-site *aak-1(nar4)* mutation. (H-J) F1 cross-progeny of *oma-1/+; oma-2/+; ghd* hermaphrodites mated with control *oma-1/+; oma-2/+* males. (K-L) F1 cross progeny from reciprocal *oma-1/+; oma-2/+; ghd* crosses. **(M)** Schematic representation of TIR-1’s conserved domains. SAM, Sterile Alpha Motif. TIR, Toll and Interleukine-1 Receptor domain. Red vertical bars, EMS-generated alleles. Red horizontal bars, Deletions. **(N-O)** Clustal Omega alignments of TIR-1 orthologs showing evolutionary conservation around the *nar65* (left) and *nar42* (right) substitutions.

TIR-1/SARM1 is an adaptor protein and metabolic sensor^24^ implicated in innate immune response^25–28^, nervous system development^29^, and neurodegeneration (both ALS-induced and Wallerian)^30–33^. It is composed of autoinhibitory armadillo repeats, two Sterile Alpha Motif (SAM) domains, and a TIR domain^26,34^, with the latter two mediating protein-protein interactions with other SAM or TIR domain-containing proteins^35,36^. The TIR domain also confers NAD+ hydrolase activity to TIR-1, required for Wallerian degeneration^37^. Interestingly, our two *tir-1* alleles had conserved residues substituted in the SAM1 *(nar65[L708F]*) or TIR *(nar42[A846T]*) coding sequences (Figure 2M-O), suggesting that both domains are required for tumor suppression.

### Canonical *pmk-1*/p38 MAPK signaling is required for homeostatic inhibition of GSC proliferation

To investigate how TIR-1 may prevent tumorigenesis in the *oma-1/2* background, we performed a second RNAi screen targeting known *tir-1* interactors. Interestingly, knockdown of *pmk-1*, the ortholog of p38 mitogen-activated protein kinase (MAPK), induced a fully penetrant Ghd phenotype (Figure 3A-B; Table S2). PMK-1/p38 is therefore, like TIR-1/SARM1, required to suppress oocyte production when their final maturation is blocked. As in mammals, PMK-1/p38 activation in *C. elegans* follows a canonical NSY-1/mitogen-activated protein kinase kinase kinase 5 (MAP3K5) and SEK-1/mitogen-activated protein kinase kinase 3 (MAP2K3) phosphorylation cascade^38,39^. Previous studies have also demonstrated that TIR-1 functions upstream of *pmk-1*/p38 signaling in the *C. elegans* innate immune response^25–28^ and neurodegeneration^32,33^. We therefore investigated if *nsy-1* and *sek-1* were also required for germline homeostasis. We found that *nsy-1(RNAi)* and *sek-1* loss-of-function each led to differentiated germline tumors in the *oma-1/2* background (Figure 3C-D, Table S2), suggesting that TIR-1 may work through canonical *pmk-1*/p38 signaling to prevent their formation.

**Figure 3.**
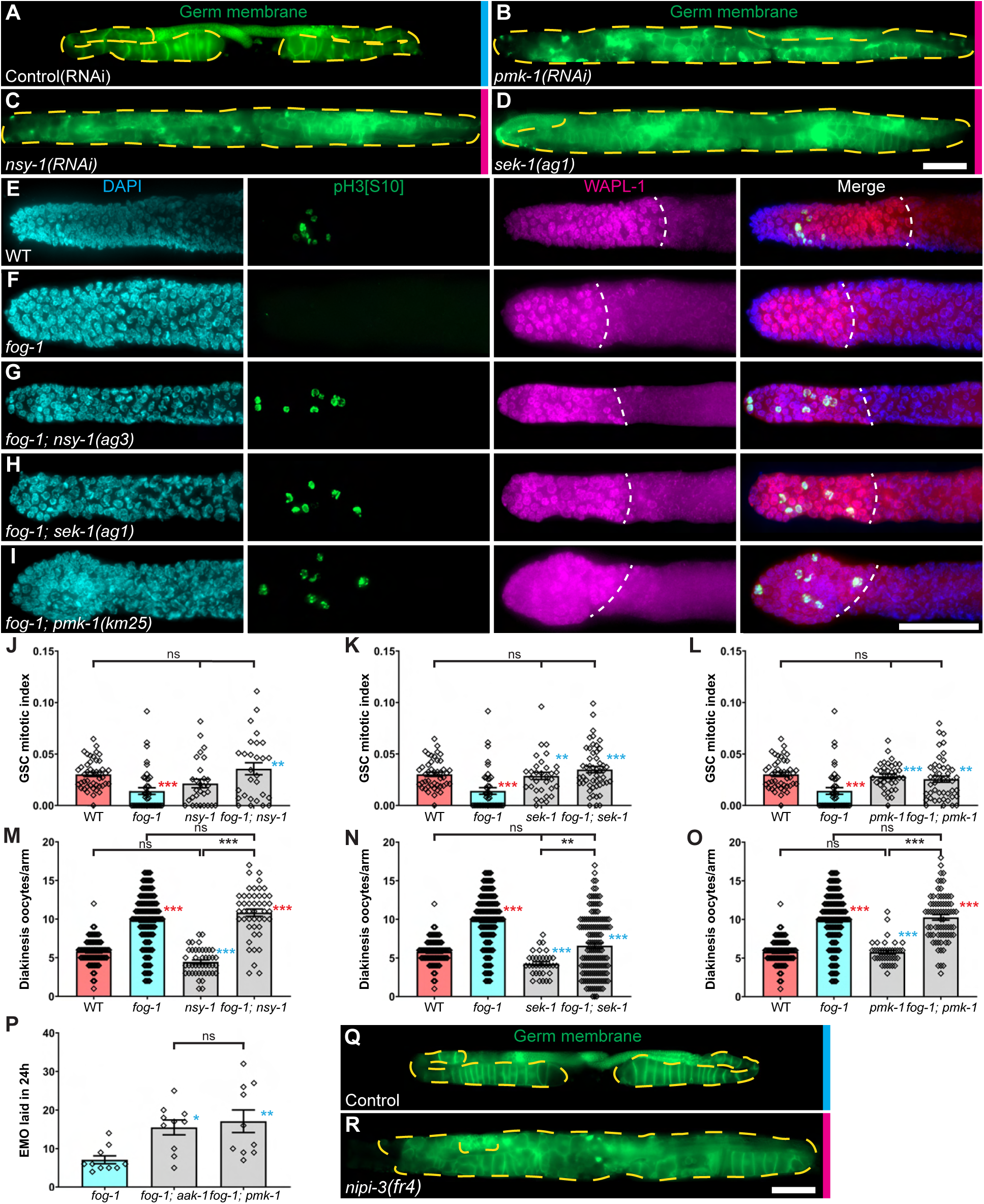
Germline homeostasis requires the canonical PMK-1/p38 signaling cascade and the NIPI-3/TRIB1 pseudokinase. **(A-D)** Representative epifluorescence micrographs of A3 hermaphrodites (A-C: 25°C; D: 20°C) of the indicated genotypes, all in the *oma-1; oma-2* background (sample sizes: 12-5-11-14). Yellow dashed lines delineate area occupied by oocytes. Anterior, left; dorsal, up. Scale bar, 100µm. A quantification of the phenotype’s penetrance is shown as a colored vertical bar on the right (Blue, Oma; Red, Ghd). All strains contain *cpSi42[Pmex-5::mNG::PLCδ-PH::tbb-2 3’UTR + unc-119(+)]*, *oma-1(zu405te33), and oma-2(te51).* **(E-I)** Representative epifluorescence micrographs of distal germlines dissected from A1 (25°C) hermaphrodites of the indicated genotypes, stained with DAPI (DNA; blue), anti-phospho[S10] histone H3 (M-phase cells^89^; green) and anti-WAPL-1 (proliferative cells^86,87^; magenta). Distal is left. **(J-L)** Average GSC MIs for each of the indicated genotype (sample sizes: (J) 45-41-29-28; (K) 45-41-32-51; (L) 45-41-37-46). **(M-O)** Average diakinesis oocyte per gonad arm for each of the indicated genotype (sample size: (M) 76-134-44-47; (N) 76-134-34-141; (O) 76-134-38-74). **(P)** Average number of EMO = endomitotic oocytes, laid by hermaphrodites between A1 and A2 stages (25°C, sample size: 10-10-10). (J-P) Each bar represents the mean ± SEM, and each point represents (J-O) a different gonad arm, or (P) the total number of EMO laid by a single adult. Red asterisks show statistical significance (p<0.05) vs wild-type; blue vs *fog-1* (Kruskal-Wallis followed by Dunn’s multiple comparisons). Double asterisks, p<0.01; triple asterisk, p<0.001. Ns, not significant. Alleles: *fog-1(q253)*, *nsy-1(ag3)*, *pmk-1(km25)*, *sek-1(ag1)*. **(Q-R)** Representative epifluorescence micrographs of A3 hermaphrodites (20°C) of the indicated genotypes, all in the *oma-1; oma-2* background (sample sizes: 14-13). Germ membranes are marked in green by *cpSi42*. Presented as in (A-D).

To confirm that *pmk-1*/p38 signaling is required for homeostatic downregulation of GSC proliferation, we crossed *nsy-1*, *sek-1,* and *pmk-1* loss-of-function alleles in a feminized *fog-1* background. As within *oma-1/2* double mutants, these animals accumulate oocytes before GSC proliferation and differentiation are inhibited^8,40^. All three mutations prevented the repression of GSC proliferation downregulation in the *fog-1* background, resulting in normal adult mitotic indexes (MIs) (Figure 3E-L). Interestingly, unlike *fog-1; aak-1* or *fog-1; daf-18* doubles, but similar to *fog-1; let-60(gf)* doubles^10^, *fog-1; nsy-1* and *fog-1; pmk-1* feminized doubles accumulated diakinesis oocytes, whereas *fog-1; sek-1* was intermediate (Figures 3M-O, S1A-E). Yet, the *fog-1; pmk-1* double mutants laid significantly more unfertilized oocytes compared to *fog-1* singles, paralleling *fog-1; aak-1* homeostasis-defective doubles, and consistent with their ongoing GSC proliferation and differentiation (Figure 3P). These results indicate that, unlike *aak-1* or *daf-18*, which also impact on ovulation control, defects in *pmk-1*/p38 signaling may more specifically prevent homeostatic repression of GSC proliferation, like let*-60(gf)*. We however note that *sek-1* may be more pleiotropic.

Consistent with its incompletely penetrant Ghd phenotype, and despite an upward trend in the GSC MI, *tir-1* loss-of-function did not significantly disrupt homeostatic induction of GSC quiescence in neither a feminized, nor an *oma-1/2* background (Figure S2A-B). Like *pmk-1*/p38 signaling mutants, the loss of *tir-1* did not prevent diakinesis oocyte accumulation (Figure S2C). These results support a role for *tir-1* upstream of *pmk-1*/p38 signaling in GSC homeostasis regulation, but its partial requirement imply the existence of additional upstream factors.

As the *C. elegans* ortholog of Tribbles homolog 1 (TRIB1) *nipi-3* was reported to activate *pmk-1*/p38 signaling in innate immunity^41,42^, we asked if was required for germline homeostasis. Consistent with this, a conditional loss-of-function allele of *nipi-3* led to a fully penetrant Ghd phenotype in the *oma-1; oma-2* background under semi-permissive conditions (Figure 3Q-R). Even under restrictive conditions however, this allele did not impair feminization-induced downregulation of GSC proliferation (Figures S2A), although feminized *nipi-3* doubles showed intermediate diakinesis oocyte accumulation (Figure S2C). As they had similar partial roles, we tested the interaction between *tir-1* and *nipi-3*. In feminized hermaphrodites, the simultaneous loss of these factors did not perturb diakinesis-stage oocyte accumulation, nor homeostatic downregulation of GSC proliferation. (Figure S2A, C). The absence of any additive or synergistic effects between *tir-1* and *nipi-3* favors the possibility that they contribute to *pmk-1*/p38 signaling activation sequentially. Altogether, our findings indicate that homeostatic repression of GSC proliferation depends on canonical *pmk-1*/p38 signaling, while NIPI-3/TRIB1 and TIR-1/SARM1 may jointly contribute to its activation.

### PMK-1/p38 signaling inhibits gonadal sheath MPK-1 to induce GSC quiescence

Next, we asked in which tissue *pmk-1*/p38 signaling acts to promote GSC quiescence in feminized hermaphrodites. An endogenous tagged *pmk-1::GFP* allele was reportedly expressed in the neurons, intestine, hypodermis and vulva precursor cells, but not in the germline^43^. Using this allele, we detected additional and relatively strong expression in the gonadal sheath nuclei, as well as in the spermathecae (Figure 4A), while feminization did not perturb this pattern (Figure 4B). We too failed to detect any PMK-1::GFP in the germline (Figure S3A-D).

**Figure 4.**
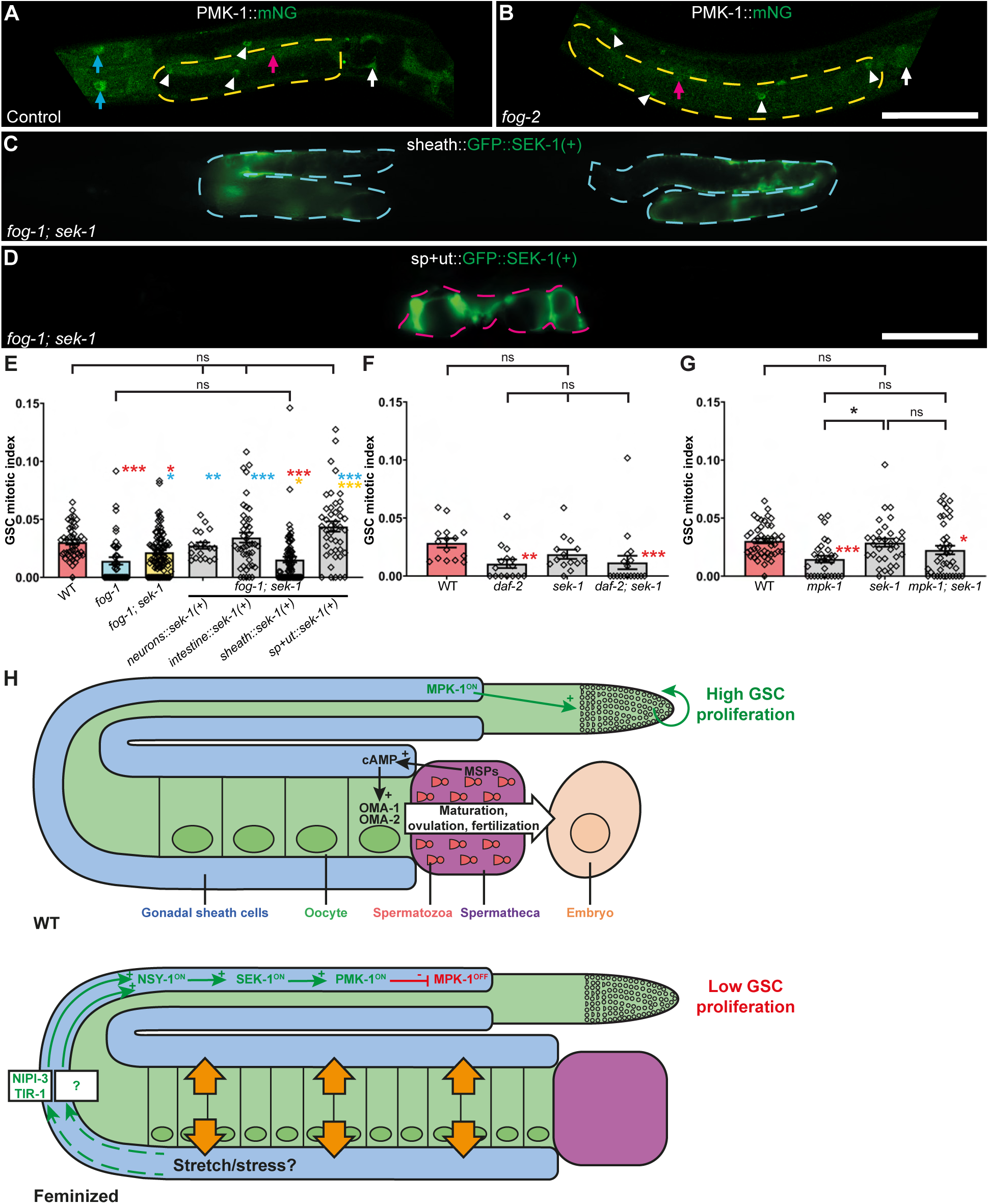
PMK-1/p38 signaling non-autonomously promotes GSC quiescence from the gonadal sheath, upstream of MPK-1/ERK. **(A-B)** Representative confocal micrographs of A1 hermaphrodites (25°C) for each of the indicated genotypes. Anterior is to the left and dorsal up. White triangles indicate gonadal sheath cells nuclei. Arrows indicate other somatic structures: blue, intestine; red, hypodermis; white, spermatheca. Yellow dashed lines delineate areas occupied by oocytes. Scale bar, 50µm. Both strains carry the *pmk-1(re170[pmk-1::mNG::3xFlag])* fluorescent reporter. **(C)** Hermaphrodites were injected with a *Plim-7::GFP::SEK-1(+)* rescue transgene. Blue dashed lines delineate gonadal sheath cells. **(D)** Hermaphrodites were injected with a *Pfos-1::GFP::SEK-1(+)* rescue transgene. Red dashed line delineates sp+ut = spermatheca and uterus. Anterior, left; dorsal, up. Scale bars, 100 µm. **(E-G)** Average GSC MIs for each of the indicated genotype (sample sizes: (E) 45-41-97-19-43-76-44; (F) 15-15-14-18; (G) 45-30-32-39). Each bar represents the mean ± SEM, and each point represents an individual gonad arm. Red asterisks show statistical significance (p<0.05) vs wild-type; blue vs *fog-1*; yellow vs *fog-1; sek-1* (Kruskal-Wallis followed by Dunn’s multiple comparisons). Double asterisks, p<0.01; triple asterisk, p<0.001. Ns, not significant. Alleles: *daf-2(e1370)*, *fog-1(q253)*, *fog-2(oz40)*, *mpk-1(ga117)*, (E) *sek-1(km4)*, (F-G) *sek-1(ag1)*. **(H)** Proposed model for homeostatic downregulation of GSC proliferation by the canonical *pmk-1*/p38 signaling cascade. MSP, major sperm proteins. Green arrows, activation. Red bars, inhibition. Yellow arrows, mechanical force exerted on the gonadal sheath cells by the accumulating oocytes.

We also tagged endogenous TIR-1 with mNeonGreen (mNG) at its C-terminus to mark all 8 isoforms. Consistent with the literature, we observed strong intestinal and neuronal *tir-1::mNG* expression^27,33^ (Figure S3E-F). As for PMK-1::GFP, we additionally detected TIR-1::mNG in the somatic gonad, but no germline expression (Figure S3E’-F’). The *tir-1::mNG* expression pattern was again, unaffected by feminization (Figure S3G-H’). The tag did not disrupt TIR-1’s function as tagged animals generated a normal number of viable adults, unlike *tir-1(Ø)* (Figure S3I)^23^. Based on the apparent absence of TIR-1 and PMK-1 from the germline, we reasoned that *pmk-1*/p38 signaling must regulate GSC proliferation cell non-autonomously from within any somatic tissues where its components are present.

To identify this critical somatic tissue, we used extrachromosomal array transgenes^44^ to drive the expression of a wild-type *sek-1* copy downstream of various tissue-specific promoters in *fog-1; sek-1* doubles, and asked whether any would rescue homeostatic downregulation of GSC proliferation. We tested *sek-1* re-expression in the nervous system (*Punc-119*), intestine (*Pges-1*), gonadal sheath (*Plim-7*) and spermatheca/uterus (*Pfos-1*)^23,45–49^ (Figure 4C-D). Gonadal sheath-specific *sek-1(+)* expression restored GSC quiescence in *fog-1; sek-1* doubles, while expression in the nervous system, intestine or spermatheca/uterus did not (Figure 4E). Together, these results show that *sek-1*, and more broadly *pmk-1*/p38 signaling, are required in the gonadal sheath to non-autonomously ensure germline homeostasis.

We next conducted epistasis analyzes to find out how *pmk-1*/p38 signaling intersects with the two pathways stimulating GSC proliferation in *C. elegans*, IIS and homeostatic signaling^10^. Since the downregulation of IIS during early larval development promotes dauer formation^50–53^ while, to our knowledge, none of the *pmk-1*/p38 signaling component have been implicated in this developmental decision, we excluded the possibility of a role for *pmk-1*/p38 upstream of IIS. To evaluate a possible downstream role, we combined a *sek-1* loss-of-function allele with a *daf-2* conditional loss-of-function that significantly reduces the GSC MI under restrictive conditions^8^. The loss of *sek-1* however failed to restore GSC proliferation in *daf-2* mutants (Figure 4F). This result indicates that *sek-1*, and most likely that *pmk-1*/p38 signaling, is not required to suppress GSC proliferation downstream of IIS downregulation.

Since homeostatic downregulation of GSC proliferation requires the inhibition of MPK-1/ERK signaling^10^, we further examined the interaction of *pmk-1*/p38 signaling components with *mpk-1*. We crossed both a *sek-1* loss-of-function and a predicted null *nsy-1* allele in a *mpk-1(Ø)* background. As both resulting double mutants had a low GSC MI, like *mpk-1(Ø)* singles (Figure 4G, S2D), we infer that *pmk-1*/p38 signaling likely acts upstream of *mpk-1* to downregulate GSC proliferation. These results are consistent with a model in which the lack of sperm and/or oocyte accumulation would, partially through TIR-1/SARM1 and NIPI-3/TRIB1, activate canonical *pmk-1*/p38 signaling in the animal’s gonadal sheath cells, from where this would in turn, inhibit MPK-1/ERK to cell non-autonomously promote GSC quiescence.

The p38 MAPK pathway is usually activated in response to various stresses, including pathogen exposure^39^, oxidative^54^, heavy metal^55^ or mechanical stresses^56–64^. We observed that oocyte accumulation was correlated with an increased proximal gonad length in feminized animals (Figure S4A-C). Oocyte accumulation therefore mechanically stretches the proximal sheath cells. We suggest that this mechanical stretching of the gonadal sheath cells could act as an upstream stress-like signal activating *pmk-1*/p38 signaling, in turn leading to the inhibition of MPK-1/ERK and GSC proliferation (Figure 4H).

## Discussion

Several studies have investigated the role of p38 MAPK signaling on stem, progenitor, and cancer cell proliferation, but disagreed on whether it has an oncogenic or tumor-suppressive activity^65,66^. Our work reveals a clear tumor-suppressive role for *pmk-1*/p38 signaling in *C. elegans*, where it prevents GSC proliferation and differentiated germline tumor formation when sperm is absent, or oocytes are fertilization-incompetent. We showed that *pmk-1*/p38 signaling acts cell non-autonomously from the somatic gonadal sheath to promote GSC quiescence. The p38 MAPK was similarly reported to regulate SC proliferation cell non-autonomously in murine hair follicles. During the telogen phase, Caspase-3 cleaves Dusp8, a p38 MAPK inhibitor, thereby triggering a Wnt3-dependent signal that stimulates neighboring SC proliferation^67^. This conserved pathway may thus indeed have oncogenic or tumor-suppressive roles depending on context.

Few studies have explored a possible role for TIR-1/SARM1 in the regulation of cell proliferation or cancer biology. Nonetheless, SARM1 was among the most frequently mutated genes in peripheral blood samples from 60 patients with malignant ovarian germ cell tumors^68^. Additionally, SARM1 was overexpressed in three cervical cancer cell lines when compared to normal keratinocytes^69^ and its knockdown in HeLa cells led to decreased survival and increased chemosensitivity^70^. Furthermore, SARM1 physically interacts with the capsaicin receptor TRPV1 to prevent inflammatory cytokines production and maintain hepatic stellate cell quiescence^71^. Our results extend this tumor suppressive role for TIR-1/SARM1, likely through its well-characterized *pmk-1*/p38 signaling activating function^26,27^.

Although we initially identified *tir-1* through our screen, the core *pmk-1*/p38 signaling components downstream (*nsy-1*, *sek-1* and *pmk-1*) exhibited stronger defects. Their loss was indeed sufficient to restore high GSC MI in feminized animals and caused a fully penetrant, severe Ghd phenotype in the *oma-1/2* background. In contrast, TIR-1 loss resulted in an incompletely penetrant Ghd phenotype in this background and did not prevent downregulation of GSC proliferation. These results are consistent with previous studies reporting that PMK-1’s activating phosphorylation is reduced in a *tir-1* loss-of-function context but is completely abolished by the loss of *nsy-1* function^26–28^. TIR-1 thus likely cooperates with other factors to activate *pmk-1*/p38 signaling and homeostatic inhibition of GSC proliferation.

Based on its involvement as a *pmk-1*/p38 activator in the context of the innate immune response^41,42^, we asked if *nipi-3* was implicated in GSC homeostasis. The reduction of *nipi-3* function caused a slightly more severe Ghd tumorous phenotype than the loss of *tir-1*, while their combined loss was not additive. Conflicting models placed *nipi-3* either upstream of *pmk-1* activation in both *tir-1*-dependent^41^ and - independent^42,72^ pathways, as a negative regulator of *pmk-1*^73^, or suggested that *nipi-3* functions independently of *pmk-1*^74^. Our results suggest that *tir-1* and *nipi-3* both act in the same pathway and are responsible for part of the core *pmk-1*/p38 signaling module activation towards homeostatic downregulation of GSC proliferation.

In adult *C. elegans* hermaphrodites, MPK-1/ERK and IIS promote GSC proliferation in parallel^8^. Based on the absence of dauer formation defects in *pmk-1* signaling mutants and our epistasis analysis, we concluded that IIS regulates GSC proliferation independently of *pmk-1*/p38 signaling. Our results showed that canonical *pmk-1*/p38 signaling regulate GSC proliferation upstream of MPK-1/ERK activity, while in the in the context of germline homeostasis, it was required in the gonadal sheath. As MPK-1/ERK can promote GSC proliferation either from the intestine or somatic gonad^18^, the latter of which is largely composed of the sheath, we infer that *pmk-1*/p38 signaling links GSC proliferation to oocyte needs by cell-autonomously repressing *mpk-1* activity in the gonadal sheath, thereby limiting its ability to non-autonomously promote GSC proliferation from there (Figure 4H).

A cell-autonomous inhibitory role for p38 on ERK’s oncogenic activity appears conserved. In a chicken allantoic membrane model, inhibition of p38, either pharmacologically or through a dominant negative allele, led to increased ERK activating phosphorylation in D-HEp3 cells, and interrupted their dormancy^75^. Similarly, in HT1080 fibrosarcoma and PC3 cell lines, the phospho-p38 to phospho-ERK ratio inversely correlated with proliferation, while shifting this balance was sufficient to favor growth or dormancy^76^. The precise mechanism by which p38 inhibits ERK signaling however remains unclear.

Exactly how *pmk-1*/p38 signaling gets activated in the context of GSC homeostasis remain to be elucidated. In mammals, p38 activation has been widely observed in response to mechanical stretching both in muscular^56,59–63^ and stromal cells^57,64^. This mechanosensitive activation appears at least conserved throughout vertebrates, as rainbow trout cardiac fibroblasts also activate p38 upon mechanical stretch^77^. Based on this, we propose that in feminized *C. elegans*, the accumulation of large, differentiated oocytes may stretch the gonadal sheath cells that enwrap them, effectively mechanically stressing them, resulting in the activation of PMK-1 signaling specifically in this tissue.

A recent *C. elegans* study demonstrated that intestinal PMK-1 is required for the apoptotic elimination of germ cells with DNA damage, thereby preventing heritable aneuploidy^78^. Our findings therefore present a second non-autonomous role for somatic PMK-1/p38 signaling on the germline. More broadly, we expect p38 activation, whether triggered by innate immune response or other stresses, to negatively regulate SC proliferation in many other contexts. If the molecular mechanisms we described here in *C. elegans* are evolutionary conserved, we would speculate that stresses that activate p38 signaling, like contracting an infection for example, may have negative consequences on the proliferation of certain SC populations. A shift from growth to defense could indeed represent an adaptive strategy to prioritize defense/survival over growth/reproduction.

## Acknowledgments

We thank Monique Zetka for reagents. Some strains were provided by the *Caenorhabiditis* Genetics Center, which is funded by NIH Office of Research Infrastructure Programs (P40 OD010440). We are grateful to the *C. elegans* gene knockout consortium and to WormBase for their essential roles in *C. elegans* research. Research in the Narbonne laboratory is funded by grants from the NSERC (RGPIN-2019-06863, RGPAS-2019-00017, DGECR-2019-00326), the CIHR (PJT-169138), and a Research Chair from the *Fondation Marcel et Rolande Gosselin* to PN. PN is a junior 2 FRQS bursary scholar (310643).

## Author contributions

Conceptualization: AC, PN

Methodology: AC, MV, PN

Validation: AC

Formal Analysis: AC, MV

Investigation: AC (epifluorescence and confocal acquisitions; genetic crosses; whole genome sequencing data analysis; complementation tests; AAK-1 and TIR-1 conserved domain annotation; TIR-1 homologs alignment; RNAi screening; pCLT1-3 design; pCLT13-14 design and construction; UTR295, 299, 570-571, 732-733 transgenic lines establishment; immuno and DAPI staining; EMO quantification, viable adult progeny quantification), MV (whole genome sequencing: pipeline design and data analysis; AAK-1 homologs alignment), BD (EMS mutagenesis; Ghd screening and mutant isolation), JR (pCLT1-3: construction; UTR377-379 transgenic lines establishment; image acquisition for Figure 2 and S3), XL (pXA1-9 design and construction), ATCZ (DAPI staining for Figures 3 and S2; image acquisition for Figure S4), LVFD (UTR621 genetic cross), AM (RNAi screening for Table S1), JD (Ghd screening), POM (Ghd screening), PN (EMS mutagenesis; Ghd screening and mutant isolation; whole genome sequencing library preparation, transgene microinjection)

Resources: PN

Writing – Original Draft: AC

Writing – Review and Editing: AC, PN

Funding acquisition: PN

## STAR methods

### Key resources table

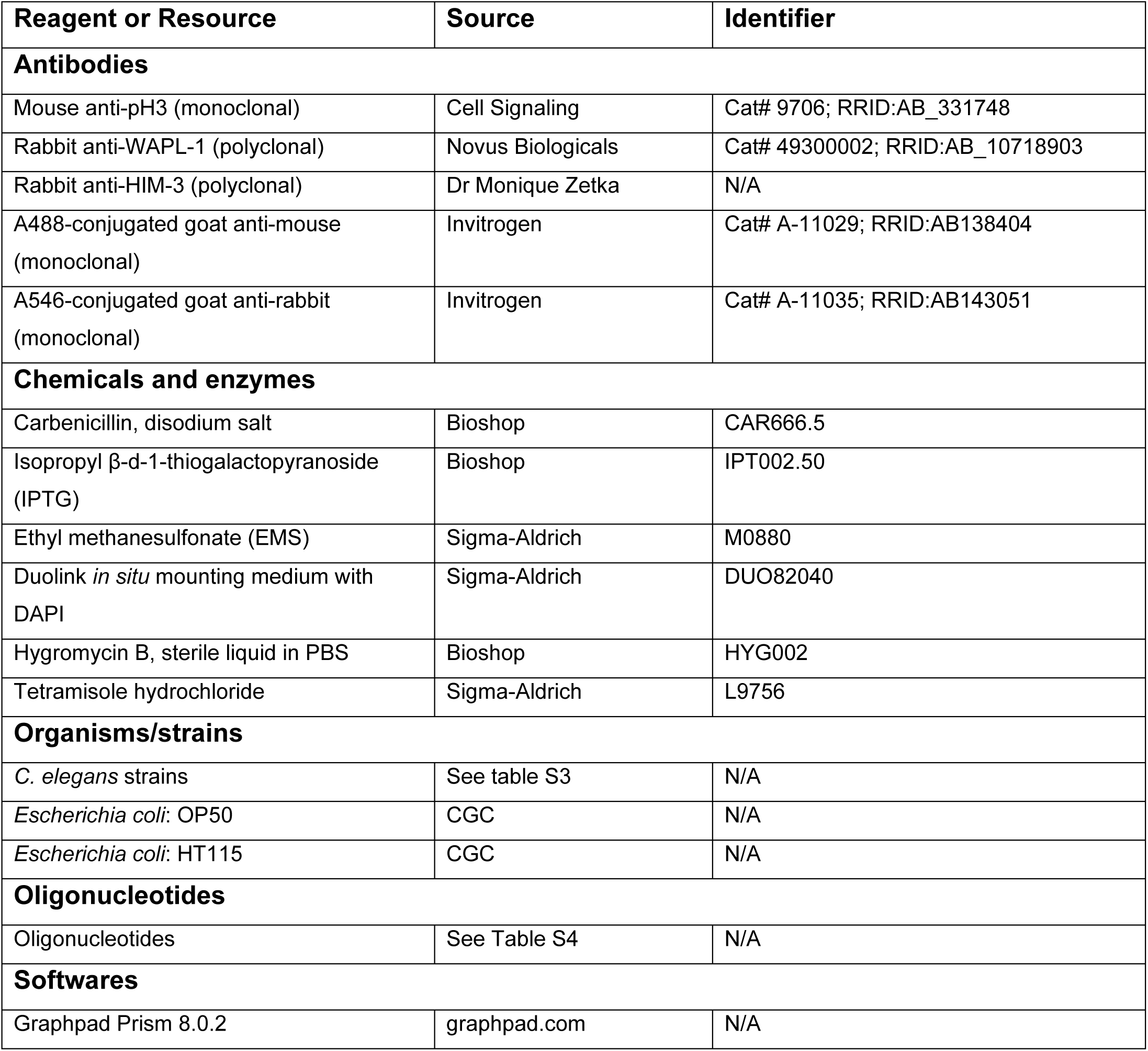
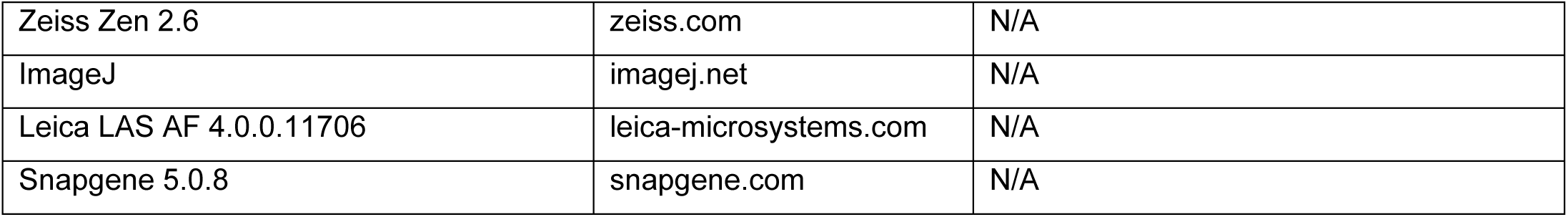

### Experimental model

#### C. elegans strains and maintenance

*C. elegans* were maintained according to standard procedures on Nematode Growth Media (NGM) plates seeded with *Escherichia coli* (OP50) at 15°C unless otherwise indicated. The Bristol isolate (N2) was used as the wild-type^21^. Table S4 lists all alleles, transgenes and rearrangements used in this study.

### Method details

#### Synchronization

Gravid adults were harvested in Milli-Q water and vortexed in a 250 mM NaOH, 1% bleach solution for approximatively 5 minutes. After 3 milli-Q water rinses, eggs were resuspended in M9 buffer and allowed to hatch for 24 hours in the absence of food under gentle agitation at 15°C. The resulting synchronized L1s were then plated on standard seeded NGM plates at 15°C unless otherwise indicated.

#### EMS mutagenesis

Synchronized *oma-1/nT1; oma-2/nT1* heteterozygous P0 late L4 larvae (complete genotype: *cpSi42[Pmex-5::mNG::PLCδ-PH::tbb-2 3’UTR + unc-119(+)]II; oma-1(zu405te33)/nT1 [qIs51] IV; oma-2(te51)/nT1 V*) were incubated for 4h in M9 containing 50 mM EMS^21^, rinsed and replated for an additional 24 hours. F1 L1s were then recovered by bleach-synchronization and plated at 15°C until they reached the L4 stage. Heterozygous *oma-1/nT1; oma-2/nT1* L4 F1s were manually transferred to new plates (5/plate) and allowed to produce F2 progeny. Once the oldest F2s had reached the L4 stage, plates were upshifted to 25°C to permit the recovery of eventual temperature sensitive alleles. Approximately 8000 haploid genomes were screened over two separate screens, and 9 independent *oma-1/nT1; oma-2/nT1* heterozygous F1s that segregated Ghd *oma-1; oma-2* animals were identified. The *oma-1/nT1; oma-2/nT1* heterozygous siblings from Ghd candidates were singled to new plates, back at 15C, to recover *oma-1/nT1; oma-2/nT1; ghd(-/-)* animals. Once homozygous, candidate mutant populations were expanded and subjected to whole-genome sequencing.

#### Whole genome sequencing and data analysis

Libraries were prepared from gDNA and sequenced by Genome Québec (Montréal, Canada) using a paired-end Illumina NovaSeq PE150. Sequencing depth varied between 40.25 M – 59.55 M (mean: 52.55 M) reads per samples.

The raw read sequences were mapped to the N2_ce11 reference genome using Bowtie2^79,80^, and processed through a variant calling and filtering pipeline, composed of MiModD^81^, SnpEff eff^82^ and SpSift Filter^83^ on the Galaxy platform^84^. Candidate *ghd* mutations were homozygous WT (+/+) in the initial strain and homozygous mutant (-/-) in the *ghd* candidates. Since candidates carried the nT1 reciprocal translocation to balance *oma-1* and *oma-2*, we also considered the variants that were heterozygous (+/-) within the nT1-balanced region (ChrIV 3,618,259-17,493,829; ChrV 1-15,080,152) in the *ghd* mutants. The *ghd-1* (*aak-1*) and *ghd-2* (*tir-1*) mutations were subsequently confirmed using Sanger sequencing.

#### Viable adult progeny quantification

Late L4-stage larvae were singled at 25°C and transferred every 24h to a new plate until individuals laid no eggs over the next 24h or ruptured due to an Egl phenotype. The number of viable adult progeny was scored.

#### GSC MI quantification

GSC MIs were evaluated as previously described^8^. As we used a temperature-sensitive *fog-1(q253)* allele, P0s were upshifted from 15°C to 25°C to prevent sperm formation in their F1 progeny. Late L4-stage F1 progeny were identified based on vulva development^85^ and picked to a new plate at 25°C, allowing them to grow for an additional 24 hours. Resulting adult day-1 (A1) animals were then harvested, and their gonads were dissected as described^10^ in a drop of PBS on a coverslip, which was then flipped against a poly-L-lysine coated slide and submitted to a freeze-crack procedure. This manipulation was always accomplished within 5 minutes^17^.

Gonads were fixed in −20C Methanol for 1 minute followed by a postfixation in 3.7% paraformaldehyde PBST (1x PBS + 0.1% Tween 20) for 30 minutes. After 1 PBST rinse and 3 10-minutes PBST washes, samples were blocked for 1 hour at room temperature in PBST containing 3% bovine serum albumin (BSA). After the blocking solution was removed, samples were incubated overnight at 4°C in primary antibody solution diluted in PBST containing 1% BSA. Anti-WAPL-1^86,87^ or Anti-HIM-3^88^ antibodies (1:500 each) were used to identify PZ cells (either WAPL-1 positive or HIM3-negative), and anti-phospho[S10]-histone H3 were used to label M-phase nuclei^89^ (1:250). After 1 PBST rinse and 3 10-minute PBST washes, gonads were incubated in the dark for 90 minutes in goat-anti-rabbit alexa546 (1:500) and goat-anti-mouse alexa488 (1:500) secondary antibody solution diluted in PBST containing 1% BSA. After 1 PBST rinse and 2 10-minute PBST washes, samples were incubated for 7 more minutes in 4’,6-diamidino-2-phenylindole (DAPI, 1µg/ml in PBS). After a final PBST rinse and a 10-minute PBST wash, gonads were mounted in 8µl Duolink *in situ* mounting medium, sealed with nail polish, and stored at −20°C until imaged. Undifferentiated germ nuclei counting in 3 dimensions was partially automated as described^90^, using an ImageJ plugin developed by Dr Jane Hubbard’s laboratory.

For *daf-2(e1370)* mutants^91^ (Figure 4F), animals were grown at 15°C until the late L4-stage to prevent dauer formation, before they were picked to a new plate and upshifted to 25°C for an additional 24h. Resulting A1 hermaphrodites were dissected, stained, imaged and analyzed as above.

#### Diakinesis oocyte quantification

Staged A1 25°C-raised F1 hermaphrodites were generated as above, fixed in Carnoy’s solution and stained with DAPI as previously described^20^. Briefly, after fixation solution removal and 3 5-minute PBST washes, samples were incubated 30 minutes at room temperature in the dark in a DAPI (0.1mg/ml in PBST) solution. After a final 5-minute PBST wash, samples were mounted in 3µl Duolink *in situ* mounting medium, sealed with nail polish and stored at −20°C until imaged. The number of diakinesis-stage oocytes (having at least one bivalent chromosome^92^) per gonad arm was then quantified. Anovulated oocytes with fully condensed chromosomes that were proximal to a diakinesis-stage oocyte were included in the counts.

#### Complementation tests

Heterozygous *oma-1/nT1; oma-2/nT1; ghd(-/-)* hermaphrodites were allowed to mate for 24h at 20°C, either with control *oma-1/nT1; oma-2/nT1*, or another *oma-1/nT1; oma-2/nT1; ghd(-/-)* candidate mutant males. Individual fertilized hermaphrodites were picked to separate plates. The resulting late L4-stage *oma-1; oma-2* homozygous F1 hermaphrodites, from plates that displayed approximatively 50% F1 males, were isolated on a new plate and allowed to grow for an additional 72h at 20°C before the Ghd phenotype was scored.

#### Image acquisition and processing

For live epifluorescence imaging (Figures 1E-P, 2A-L, 3A-D, 3Q-R and 4C-D), worms were paralyzed by soaking in a 0.1% tetramisole M9 solution and flipped onto a 3% agarose M9 pad positioned on a microscope slide. Images were acquired on an inverted Zeiss Axio Observer.Z1 using a 20x/0.8 numerical aperture dry objective, with a 1 µm z-step. For epifluorescence, samples were excited by a LED module at 488nm (50% intensity, 100ms exposure) and emission was collected at 500-550nm. Images were stitched using the Zen 2.6 software. Brightness/contrast were uniformly adjusted, and animals were computationally straightened using the ImageJ software for display purposes. All displayed live epifluorescence micrographs show single focal planes.

For fixed epifluorescence imaging (Figures 3E-I and S1A-E), samples were acquired on an inverted Zeiss Axio Observer.Z1 using a 40x/1.4 numerical aperture oil objective with a 1 µm z-step. Samples were excited by a LED module at 353nm (DAPI, 50% intensity, 50ms exposure), 493nm (alexa488, 50% intensity, 50ms exposure) or 557nm (alexa546, 50% intensity, 300ms exposure) and emissions were collected at 420-470nm, 500-550nm and 570-640nm respectively. Images were deconvolved using the Zen 2.6 software Brightness/contrast were uniformly adjusted on ImageJ for display purposes. All displayed fixed epifluorescence micrographs are maximal intensity z-projections.

For live confocal imaging (Figures 4A-B, S3A-H’ and S4A-B), hermaphrodites were paralyzed as described above. Slides were sealed with VALAP (1:1:1 vaseline, lanolin, paraffin) and images were acquired within 30 minutes using an inverted Leica SP8 point scanning confocal microscope with a 40x/1.30 numerical aperture oil objective at 0.30 µm z-steps.

For PMK-1::mNG (Figures 4A-B and S3A-D), samples were excited by a 488nm laser (5% intensity) and emission was collected at 507-527nm. Transmitted white light was collected using the same excitation with a PMT trans photoreceptor. For TIR-1::mNG (Figure S3E-H’), samples were excited by a 488nm laser (2% intensity) and emission was collected at 507-528nm. For proximal gonad length measurements (Figure S4A-B), samples were sequentially acquired using a 405nm (0.5% intensity) laser excitation with a 413-463nm emission collection, and a 552nm laser excitation (0.5% intensity) laser excitation with a 574-594nm emission collection. A segmented line from the gonad arm bend to the distal spermatheca, going through the center of each oocyte, was drawn and measured using ImageJ. Only anterior gonad arms were analyzed.

To reduce noise and amplify signal in all confocal acquisitions, each pixel was acquired three times, and the intensity values were averaged by the Las AF software. Brightness/contrast were uniformly adjusted for display purposes using ImageJ. Displayed confocal micrographs are either single focal planes (Figures 4A-B, S3A-D and S3E’-H’) or maximal intensity projections (Figures S3E-H and S4A-B). To ensure signal consistency between individuals and samples, only gonad arms located above the intestine were considered.

#### RNA interference

RNAi screens were conducted as previously described^93,94^. Briefly, L4-staged *oma-1/nT1; oma-2/nT1* P0s were picked to 25°C NGM plates supplemented with 25µg/ml carbenicillin and 1mM IPTG and seeded with *E. coli* HT115 either containing an empty L4440 vector or expressing an exon-rich dsRNA sequence targeting a gene of interest. The resulting late L4-staged *oma-1; oma-2* homozygous F1s were isolated on a new plate and allowed to grow for an additional 72h at 25°C before their phenotype was scored. RNAi constructs were obtained from the Ahringer library^93^ except for the *nsy-1*, *pmk-1* and *pmk-3* RNAi clones, which were generated as described below. All clones were validated by Sanger sequencing.

#### Plasmids, transgenesis and CRISPR/Cas9-mediated genome edition

We used the Gibson *et al.*, 2009 protocol^95^ for all plasmid assembly, except for pCLT2-3 and pCLT13, which were generated using kinase ligase DpnI (KLD) site-directed mutagenesis^96,97^ on the pDD162^98^ or pXA1 vectors, respectively. The source DNA and primers that were used to generate all plasmids as well as their microinjection concentrations are indicated in Table S3. Extra-chromosomal arrays were generated as previously described^44^ by regular plasmid DNA microinjections at a total concentration of 200 µg/ml using a pKSII as a filler DNA and pCFJ104[Pmyo-3::mCherry]^99^ (5 µg/ml) as a co-injection marker.

TIR-1::mNG fluorescent knockins were generated as previously described^100^. A pCLT1 repair template (self-excising cassette (SEC)::mNG::3xFLAG-TIR-1 homology arms, 10 µg/ml) was co-injected with pCLT2 and pCLT3 constructs (sgRNA-Cas9, each at 25 µg/ml) and pCFJ104[Pmyo-3::mCherry] co-injection marker (5 µg/ml). Following hygromycin (250µg/ml) selection, mCherry negative Rol F1s were isolated on individual plates. F2s were isolated from F1s that segregated 100% Rol progeny. Three independent F2 edited lines were selected and their L1/L2-staged F3 progeny was heat shocked at 34°C for 4 hours for SEC removal.

#### Sequence alignment

Protein sequences were obtained from Wormbase^101^. Protein alignments were conducted on SnapGene using the Clustal Omega algorithm^102^. Conserved domains were obtained from UniprotKB’s protein annotation^103^.

### Quantification and statistical analyses

Graphs were generated and statistical analyses were performed using GrapPad Prism 8.0.2. For each dataset, the Shapiro-Wilk normality test was applied to determine whether it followed a Gaussian distribution. Homoscedasticity was then verified either with an F test when comparing 2 samples, or with the Brown-Forsythe test when comparing >2 samples.

When both normality and homoscedasticity were verified, a parametric test was used (unpaired t-test, when comparing 2 samples). When either normality or homoscedasticity was not verified, a non-parametric test was used (Mann-Whitney test, when comparing 2 samples or Kruskal-Wallis test followed by Dunn’s multiple comparisons when comparing >2 samples). Statistical significance was marked by asterisks as follows: one asterisk, p<0.05; two asterisks, p<0.01; and three asterisks, p<0.001.

Except for EMO (Figure 3P) and viable adult progeny (Figure S3I) quantifications, which present single replicates, all presented datasets are a merge of ≥ 2 independent replicates.

## Supplemental information

**Figure S1.**
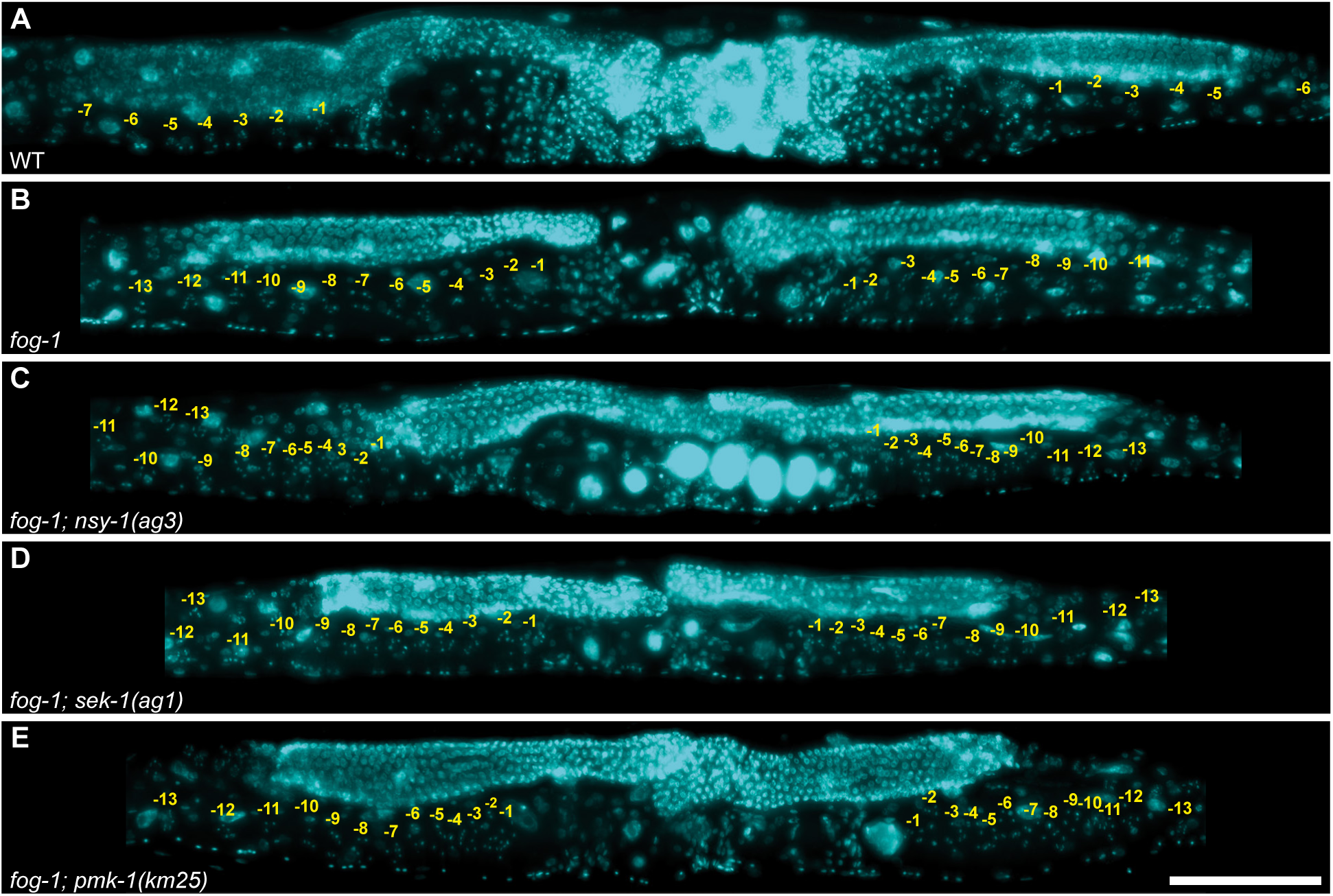
Feminized PMK-1/p38 signaling double mutants accumulate diakinesisoocyte. **(A-E)** Representative epifluorescence micrographs of A1 hermaphrodites (25°C) of the indicated genotypes, stained with DAPI (DNA; blue). Yellow numbers indicate diakinesis-staged oocytes. Anterior, left; dorsal, up. Scale bar, 100μm. Allele: *fog-1(q253)*. numbers indicate iakinesis-staged oocytes. Anterior, left; dorsal, up. Scale bar, 100µm. Allele: *fog-1(q253)*.

**Figure S2.**
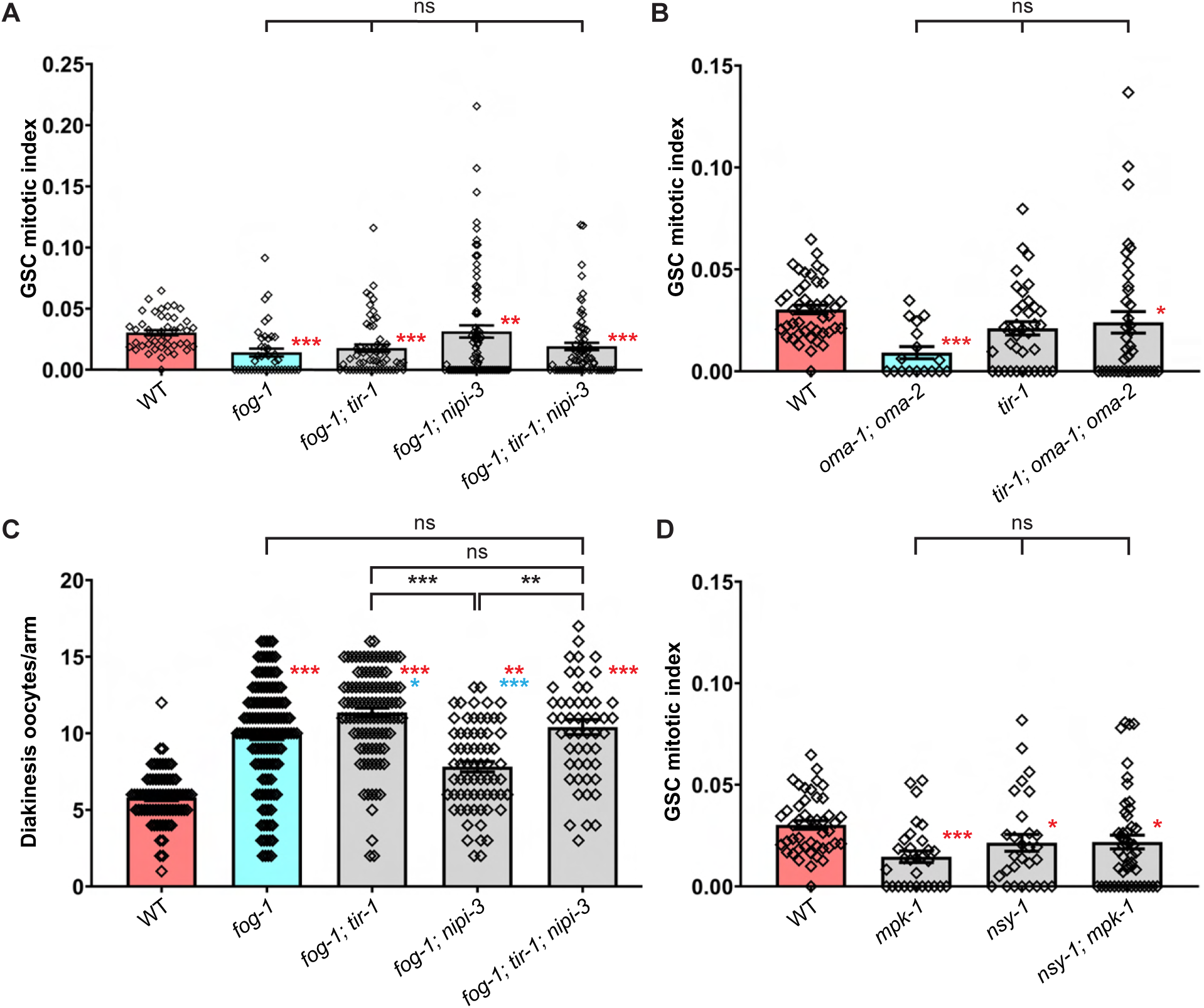
Genetic requirement for homeostatic downregulation of GSC proliferation. **(A-B)** Average GSC MIs for each of the indicated genotypes (sample sizes: (A) 45-41-58-80-73; (B) 45-17-37-38). **(C)** Average diakinesis oocyte per gonad arm for each of the indicated genotype (sample sizes: 76-134-104-67-52). (A-C) Each bar represents the mean ± SEM, and each point represents a different gonad arm. Red asterisks show statistical significance (p<0.05) vs wild-type; blue vs the appropriate *fog-1* or *oma-1; oma-2* control (Kruskal-Wallis followed by Dunn’s multiple comparisons). Double asterisks, p<0.01; triple asterisk, p<0.001. Ns, not significant. **(D)** Average GSC MIs for each of the indicated genotype (sample sizes: 45-30-29-49). Statistics and graphical representation as in (A-C). Alleles: *fog-1(q253)*, *mpk-1(ga117)*, *nipi-3(fr4)*, *nsy-1(ag3)*, *oma-1(zu405te33)*, *oma-2(te51)* and *tir-1(ok2859)*.

**Figure S3.**
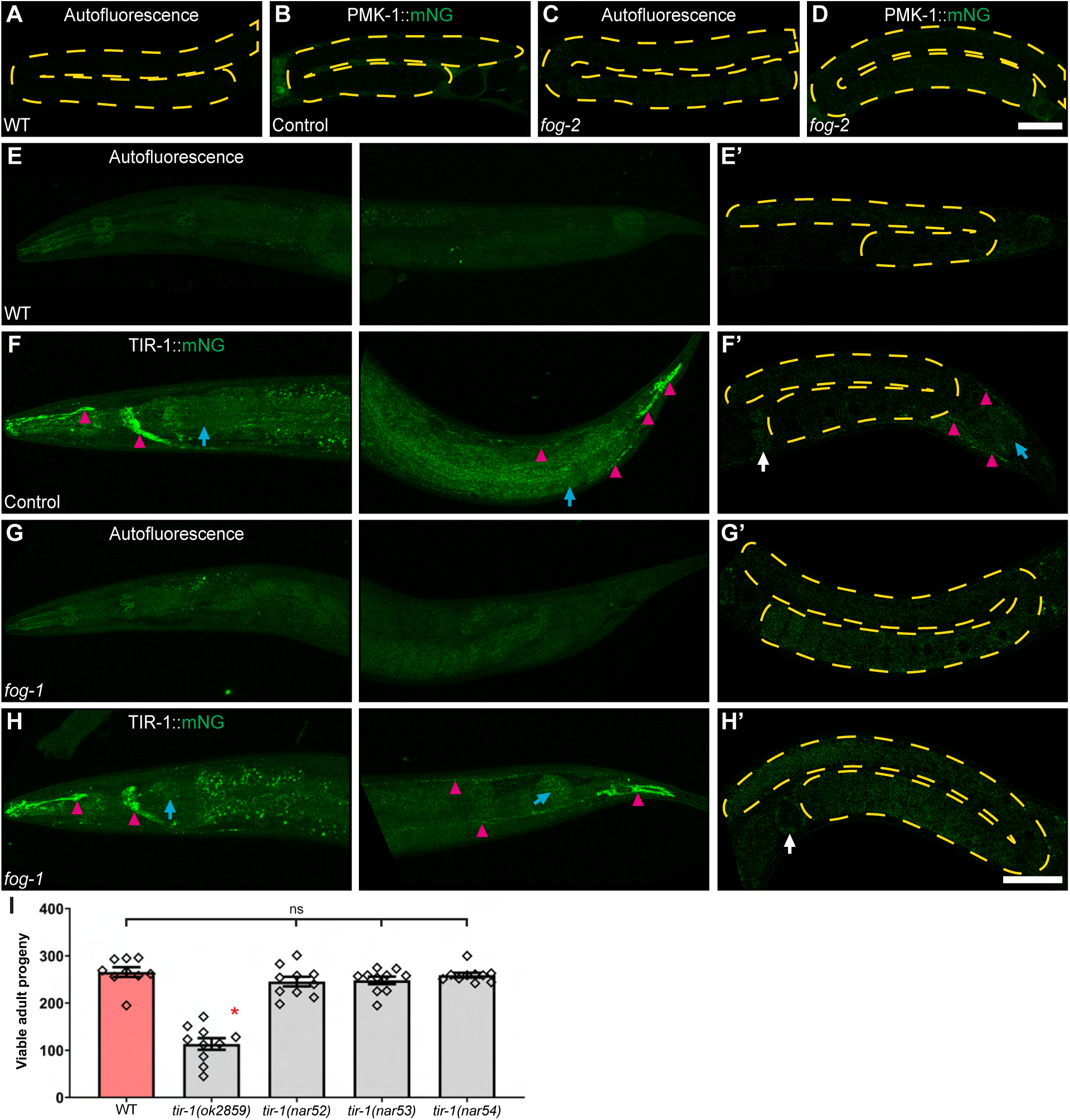
Insights on the expression patterns of PMK-1::mNG and TIR-1::mNG. **(A-H’)** Representative confocal micrographs of A1 hermaphrodites (25°C) for each of the indicated genotypes.Yellow dashed lines delineate anterior gonad arm. Red triangles indicate nervous structures. Arrows indicate other somatic structures: blue, intestine; white, spermatheca. Reporter strains carry (B, D) the *pmk-1(re170[pmk-1::mNG::3xFlag])* or (F, H) the *tir-1(nar53[tir-1::mNG::3xFlag])* fluorescent reporter. (A-H’) Anterior is to the left and dorsal up. Scale bar, 50µm. **(I)** Average viable adult progeny for each of the indicated genotypes (sample sizes: 9-10-10-10-10). Each bar represents the mean ± SEM, and each point represents the number of viable F1 adults generated by an independent P0. Red asterisk shows statistical significance (p<0.05) to all other samples (Kruskal-Wallis followed by Dunn’s multiple comparisons). ns = non-significant. Alleles: *fog-1(q253)*, *fog-2(oz40)*.

**Figure S4.**
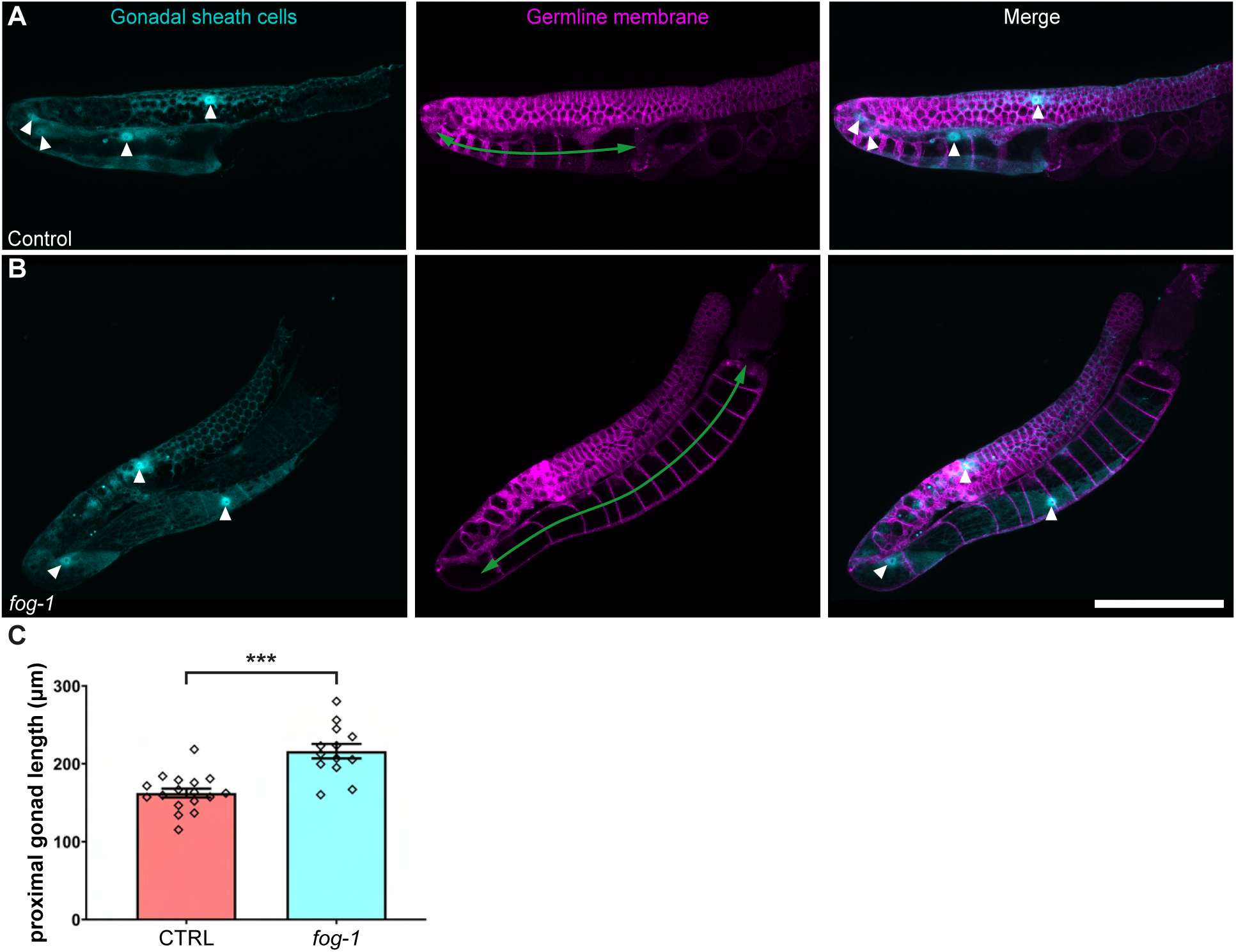
Oocyte accumulation is linked to proximal sheath cells stretching. **(A-B)** Representative confocal micrographs of A1 hermaphrodites (25°C) for each of the indicated genotypes. (A, B) gonadal sheath cells are marked in blue by narEx110. White triangles indicate gonadal sheath cells nuclei. Germ cells are labelled in red with cpSI20. Green double arrows outline the proximal gonad. Anterior, left; dorsal, up. Scale bar, 50µm. Both strains contain *narEx110[Plim-7::BFP; Pmyo-2::GFP]* and *cpSi20[Pmex-5::TAGRFPT::PH::tbb-2 3’UTR + unc-119(+)]*. **(C)** Average proximal gonad length of A1 hermaphrodites (25°C) for each of the indicated genotypes (sample sizes: 17 and 13). Each bar represents the mean ± SEM, and each point represents an individual anterior gonad arm. Triple asterisks show statistical significance between the two samples (unpaired t-test). Allele: *fog-1(q253)*.

**Table S1:** RNAi screen of multi-hit candidates identifies *tir-1* as being implicated in germline homeostatic signaling.

| RNAi-targeted genes | Mutants | Human homolog(s) | Oocyte hyperaccumulation* |
| --- | --- | --- | --- |
| *aak-1* | *nar4*, 5, 38 | PRKAA2 | + |
| *adt-1* | *nar1*, 4 | ADAMTS18 | - |
| *agef-1* | *nar1*, 2 | ARFGEF2 | - |
| *C24A3.4* | *nar4*, 37 | AMACR | - |
| *C46A5.4* | *nar1*, 36 | PXDN | - |
| *cla-1* | *nar2*, 33, 42, 43 | RIMS2 | - |
| *ctf-18* | *nar3*, 37 | CHTF18 | - |
| *dig-1* | *nar1*, 4, 35, 37 | PTPRF | - |
| *F25G6.9* | *nar33*, 43 |  | - |
| *fbxa-136* | *nar5*, 40 |  | - |
| *irld-45* | *nar3*, 39 |  | - |
| *lips-2* | *nar1*, 43 |  | - |
| *myo-5* | *nar3*, 4, 33 | MYH4 | - |
| *nas-19* | *nar1*, 3 | TLL1 | - |
| *nep-1* | *nar34*, 39 | ECE1/2, KEL | - |
| *nhr-47* | *nar4*, 36, 43 | HNF4A | - |
| *nhr-258* | *nar2*, 37 | HNF4A | - |
| *ret-1* | *nar2*, 41 | RTN1/3/4 | - |
| *T19B10.5* | *nar32*, 34 | AHNAK2 | - |
| *tir-1* | *nar65\*\**, 42 | SARM1 | + |
| *vab-7* | *nar3*, 43 | EVX1 | - |
| *W01B6.8* | *nar39*, 40 |  | - |

\*When targeted by RNAi in the *oma-1*; *oma-2* genetic background.
\*\*As the *ghd(nar4)* mutant carried alleles of two *ghd* genes, the *ghd-2/tir-1(nar4)* allele was renamed *tir-1(nar65)*.

**Table S2:**
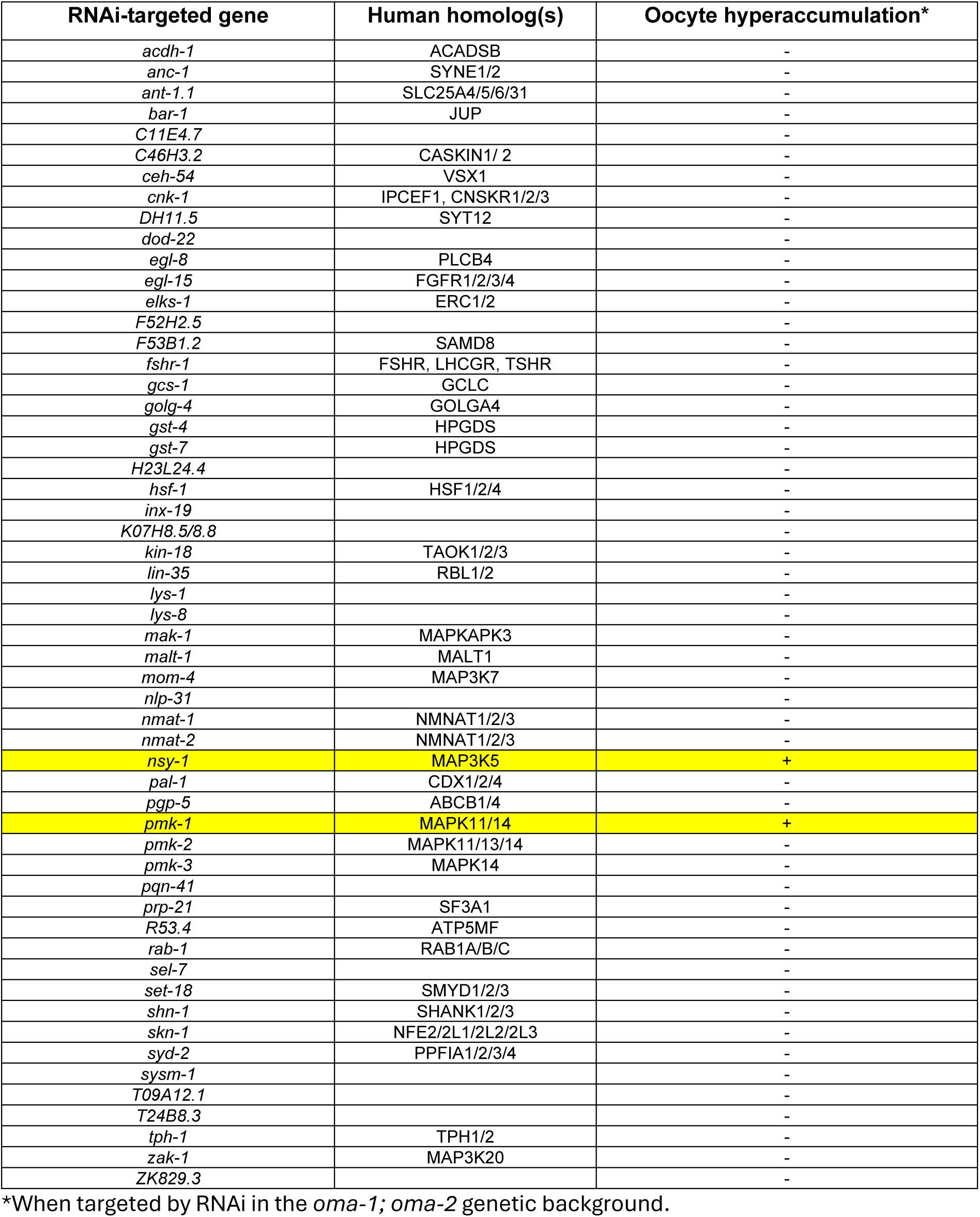
RNAi screen of *tir-1* interactors highlights *nsy-1* and *pmk-1* as required for homeostatic inhibition of GSC proliferation.

**Table S3:**
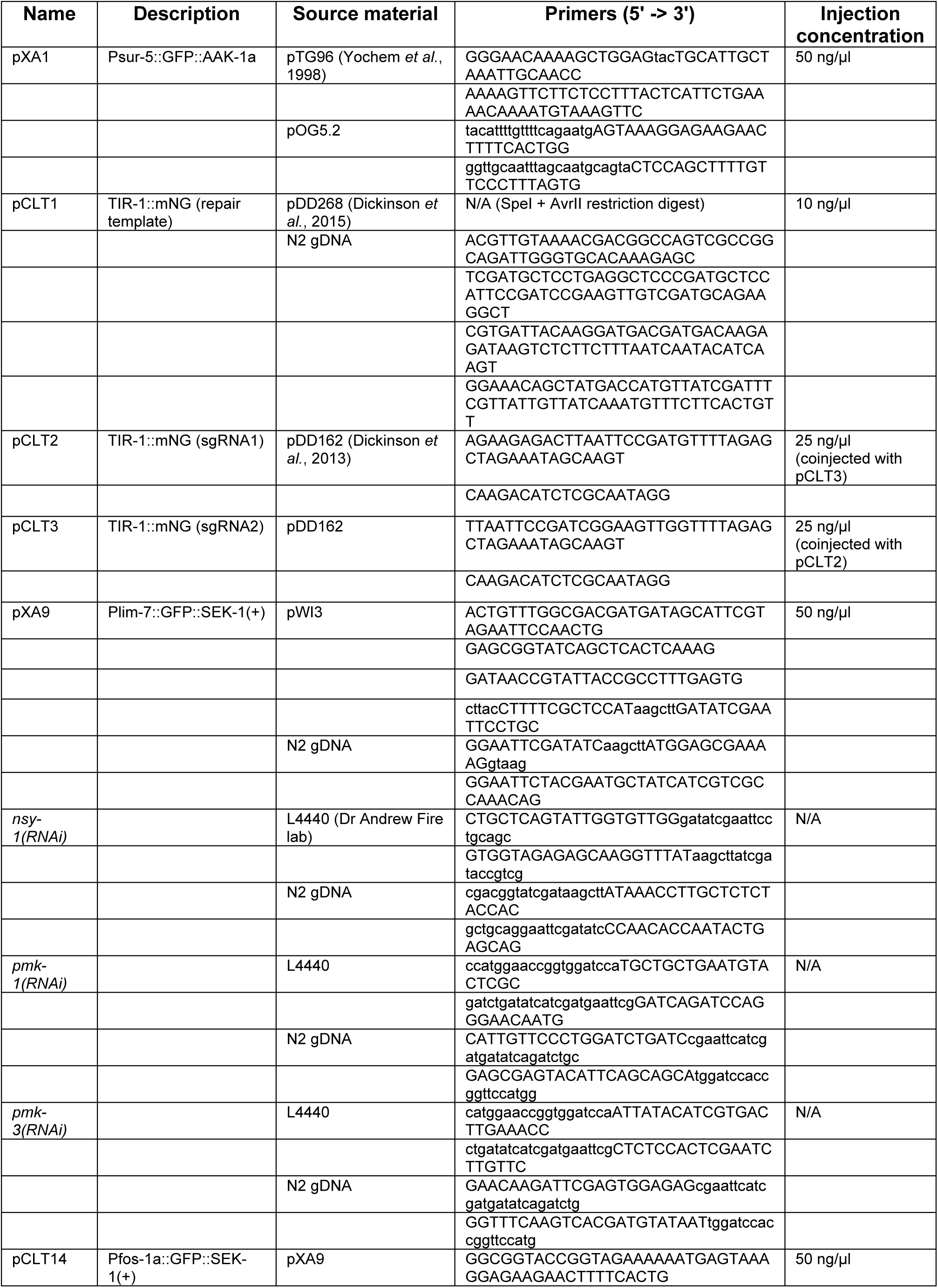
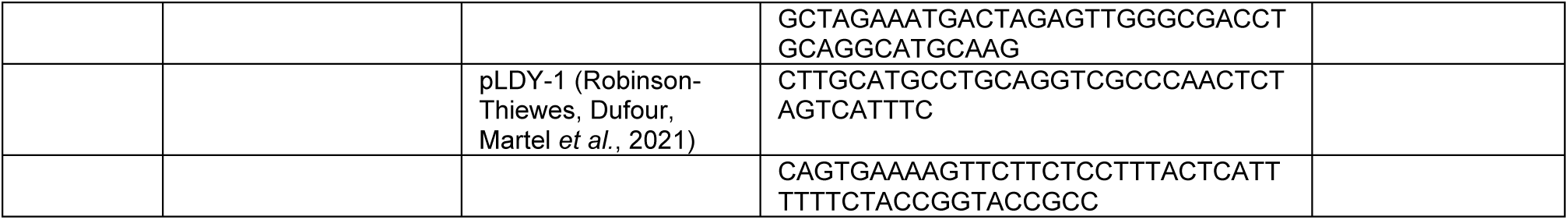
Plasmid design. Related to STAR methods.

**Table S4:**
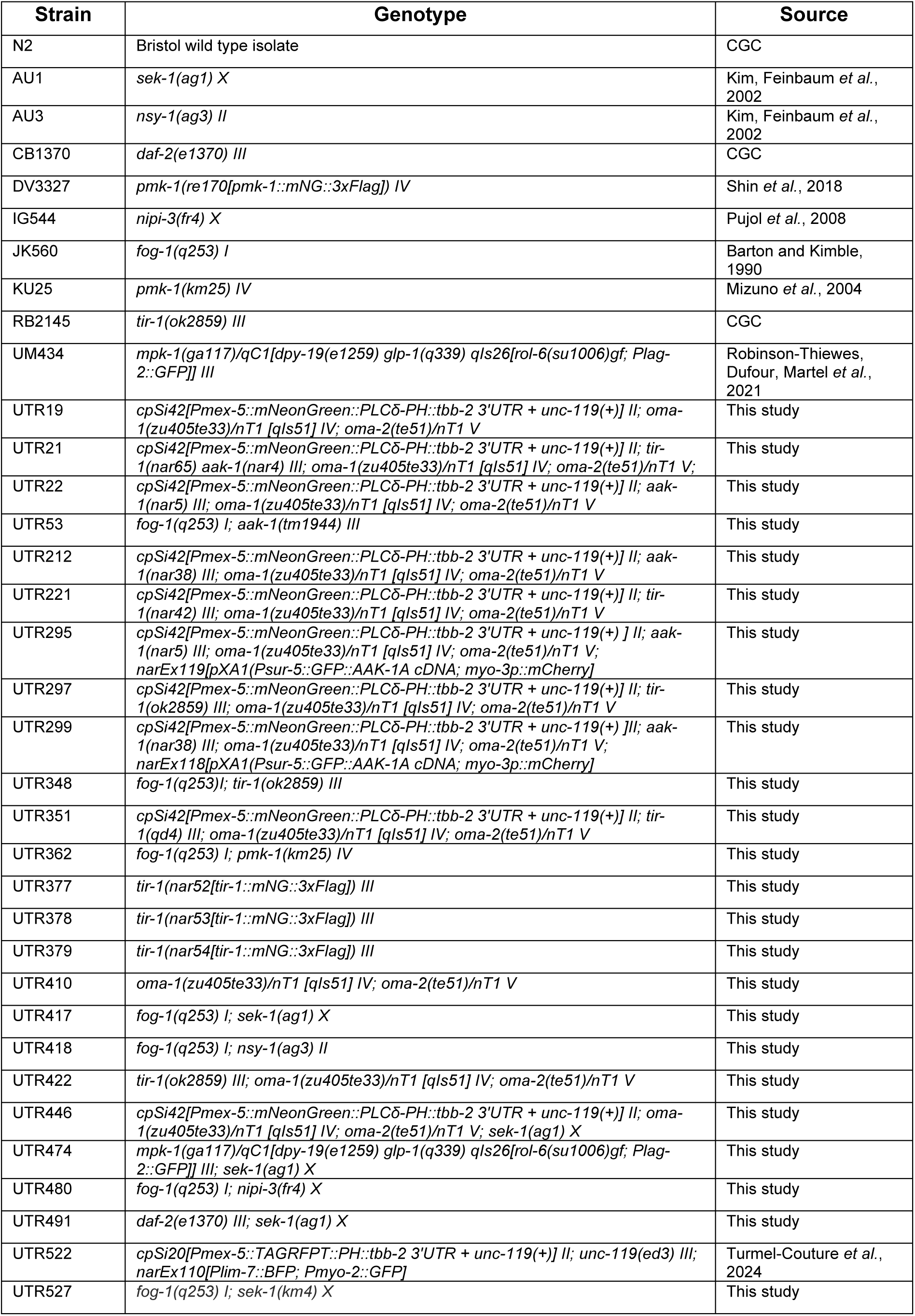
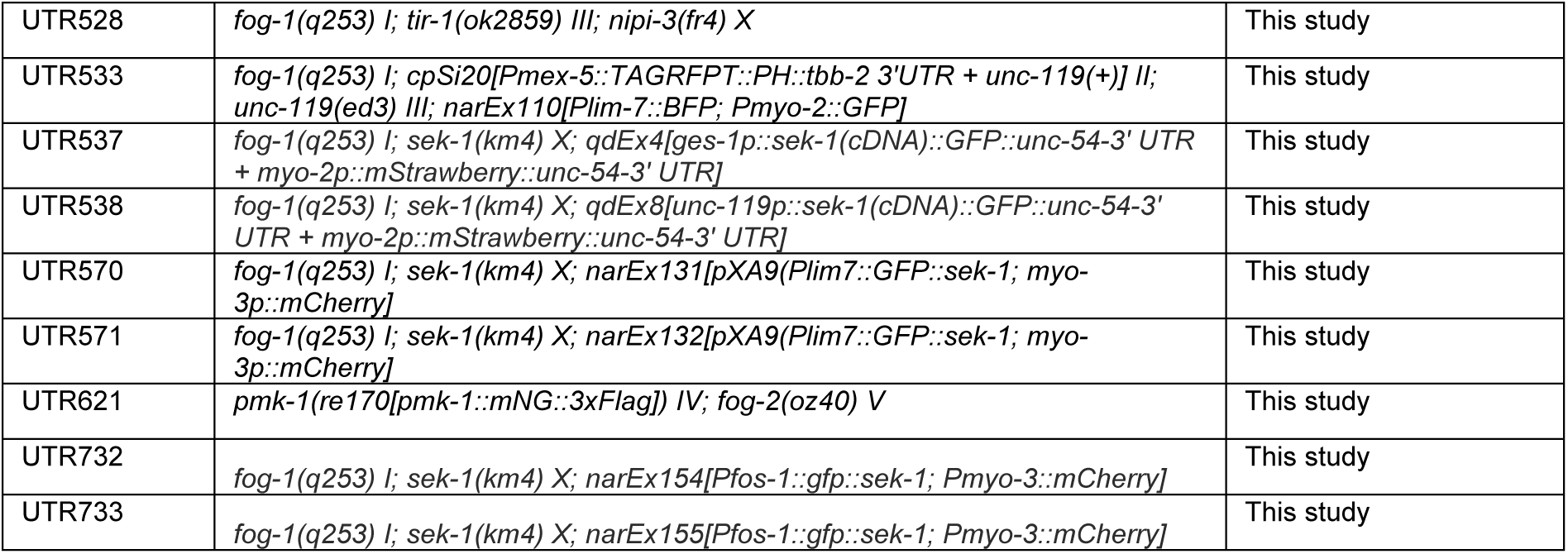
Strains, alleles, transgenes and rearrangements used in this work. Related to STAR Methods.

